# Polymer translocation through nano-pores: influence of pore and polymer deformation

**DOI:** 10.1101/313684

**Authors:** M. A. Shahzad

## Abstract

We have simulated polymer translocation across the a *α*-hemolysin nano-pore via a coarse grained computational model for both the polymer and the pore. We simulate the translocation process by allowing the protein cross a free-energy barrier from a metastable state, in the presence of thermal fluctuations. The deformation in the channel, which we model by making the radius of pore change from large to small size, can be originated by the random and non-random (systematic) cellular environment, drive out the polymer out of equilibrium during the transport dynamics. We expect that in more realistic conditions, effects originating on the translocation phenomena due to the deformability of the nano-pore can either decrease or increase the transport time of biomolecule passing through the channel. Deformation in channel can occurred because the structure of *α*-hemolysin channel is not completely immobile, hence a small pore deformation can be occurred during translocation process. We also discuss the effects of polymer deformation on the translocation process, which we achieve by varying the value of the empirical and dihedral potential constants. We investigate the dynamic and thermodynamical properties of the translocation process by revealing the statistics of translocation time as a function of the pulling inward force acting along the axis of the pore under the influence of small and large pore. We observed that a pore with small size can speed down the polymer translocation process, especially at the limit of small pulling force. A drastic increase in translocation time at the limit of low force for small pore clearly illustrate the strong interaction between the transport polymer and pore. Our results can be of fundamental importance for those experiments on DNA-RNA sorting and sequencing and drug delivery mechanism for anti-cancer therapy.

## I. INTRODUCTION

The passage of a molecule through a biological pore, the so-called translocation, is a fundamental process occurring in a variety of biological processes [1], such as virus infection [2], and gene expression in eukaryotic cells [3]. The translocation process involves the exchange of proteins, ions, energy sources, genetic information and any particle or aggregate that plays a role in cell functioning and living. For instance, translocation of bio-polymer across membranes channel is universal in biology and includes the passage of RNA strands inside nuclear pores [4], absorption of oligonucleotides on suitable sites [5], transport of protein back and forth from the cells [6], mitochondrial import, protein degradation by Adenosine triphosphate (ATP)-dependent proteases and protein synthesis [7, 8]. The transport can be either passive or active. The passive transport involves the movement of biochemicals from areas of high concentration to areas of low concentration and requires no energy for transportation. The passive transport can take place by diffusion or by interposition of carriers to which the transported substances are bounded. On the other hand, active transport are biological processes that move oxygen, water and nutrients into cells and remove waste products. Active transport requires chemical energy because it is the movement of biochemicals from areas of lower concentration to areas of higher concentration. In active transport, trans-membrane proteins use the Adenosine-Tri-Phosphate (ATP) chemical energy to drive a molecule against its concentration gradient. A mechanism of selection, based on the interaction between signal recognition particles and relative receptors, allows the trans-membrane proteins to detect the system to be transported [9].

In the recent past, translocation processes of biomolecules through nano-pores include interesting experimental and theoretical issues at the core of an intense research activities. Experimentally, the first successfully voltage-driven DNA translocation through a *α*-hemolysin channel (protein channel) embedded in lipid bilayer membranes was obtained in 1996 by Kasianowicz et al. [12]. In this device.one bio-polymer is introduced on the cis-side, it crosses the pore driven by thermal fluctuation and by the voltage and flows into the opposite chamber, the trans-side. Its passage temporarily clogs the channel and provokes a detectable ion current drop, which strongly depends on the chemical and physical properties of the molecule that occupied the pore. Moreover, the duration of current loop is a direct measure of the translocation time, which is in turn used to characterize a single transport event, along with the intensity of the blockage itself. Since then, translocation phenomena have been used to characterize single objects like single-stranded or double-stranded DNA, proteins, or even cells [11, 13–20]. In addition to biological pore, solid-state nanopores have also been used for detection and characterization of protein [21, 22], DNA/protein [23] or nano-materials/nano-particles [24–29].

Many research group has discussed the controlled unfolding and translocation of proteins through the *α*-hemolysin pore using AAA+ unfoldase ClpX [30–40]. AAA+ machine unfold and translocate polypeptide into associated peptidases. ClpX (an unfoldase) generates mechanical force to unfold its protein substrate from Escherichia coli, alone, and in complex (with combination) with the ClpP peptidase. ClpX hydrolyzes ATP to generate mechanical force and translocate polypeptide chain through its central pore [36]. AAA+ unfoldases target structurally and functionally diverse proteins in all cells. Also, AAA+ target proteins not only in a folded, soluble conformation, but also in hyperstable mis-folded or aggregated states. These molecular machines must use efficient mechanisms to unravel proteins with a wide range of thermodynamic stabilities, topologies, and sequence characteristics. These results shows that ClpX(P) is able to generate and apply mechanical forces sufficient to unfold most target proteins.

Recently, some theoretical and experimental work has been done on translocation processes where the cross-section of the pore changes during the passage of biomolecule through channel. In such translocation process, the dynamical nature of the pore enable the polymer chain to translocate more efficiently as compare to the translocation of polymer in static pore [47]. On the experimental side, it has been shown that the translocation of DNA through a nano-channel can be modulated by dynamically changing the cross-section of an elastomeric nano-channel device by applying mechanical stress [48–52]. Time-dependent driving force are also used in the translocation process [53–58]. There are some biological examples of such fluctuating environment in translocation are the nuclear pore complex, which plays an essential role in nucleocytoplasmic transport in eukaryotes [59]. Another example is the exchange of molecules between mitochondria and the rest of cell which is controlled by the twin-pore protein translocate (TIM22 complex) in the inner membrane of the mitocondria [60]. Moreover, using an alternating electric field (time-dependent driving force) in the nanopore has been implied as a source for DNA sequencing [61]. It is worth to mention an interesting results of Menais et al [64] and Fabio et al [65], where interplay between thermal fluctuations and the deformability properties of the channel play an important role in characterizing the translocation process of polymer and proteins.

Here we discuss the translocation of a polymer through a *α*-hemolysin pore. The work is based on Langevin molecular dynamics simulations exploiting the so-called Gō-like model [66, 67]. We address the issue that how the deformation of channel, and polymer affects the translocation process. We coarse-grained molecular dynamics simulations for both proteins and pores. The simulations results are interpreted by comparison using the drift-diffusion Smoluchowski equation with radiation boundary conditions to understand the phenomenology of proteins translocation. The motivations to understand the effects of fluctuating environment in protein translocation are: (a) Fluctuations of the membrane pore (b) Thermal fluctuations of environment (normal modes), (c) Fluid density variation, (d) Cycles of ATP concentration, (e) Alternating electrical potential in voltage driven experiments, (f) Systematic (non-random) effects in chemical conditions. All these elements prompted to consider translocation process across a fluctuating pore. In particular we consider translocation of bio-molecules under the influence of deformation of the channel, and polymer.

The paper is organized as follows: sec.II we describe the computational model used to simulate polymer in a confinement cylindrical geometry. We used coarse grained Gō model both the polymer and confinement geometry. In sec.III we discuss the results of polymer translocation through a dynamical nanopore. Sec. IV is devoted to the conclusion.

## II. COMPUTER MODELING OF STRUCTURED POLYMER CHAIN AND NANOPORE

We used a coarse-grained computational model to simulate numerically the translocation process for the polymer through a channel. We consider the Gō-like model as a natural approach to access the impact of the molecules structural properties on translocation. Below we show in detail the importing mechanism, pore and particle interaction potential, numerical integration scheme and equation of motion for the material points constituting the system.

### A. Polymer numerical model: The Gō-like approach

The model polymer we have considered is flexible and presents a base structure mimicking a folded protein at the coarse-grained level. A linear polymer chain is organized into a space-filling, compact, and well-defined 3-dimensional structure. The monomers in the polymer lie in the same plane, and successive planes defined angels *ϕ* and *ψ*. The conformation of a chain of *n* polymer can be defined by 2*n* par. The approach presented in our numerical work is adopted from reference [66–74, 76, 77].

Using the Gō-like approach for the numerical simulation, we used the following potentials acting on the material point of the polymer: (1) Peptide potential (or bond potential) *V*_p_, (2) Bending angle potential *V*_θ_, (3) Twisted angle potential *V_ϕ_*, (as shown in Fig. (1)) and (4) Non-bonded interactions (Lennard-Jones potential or barrier) *V_nb_*. These interactions are the simplest potential energy function that can reproduce the basic feature of protein energy landscapes at a mesoscopic level of detail, and it has proved to give insight into a remarkably broad range of properties. The combinations of a potential energy function and all the parameters that go into it constitutes a so-called force field.

**FIG. 1.**
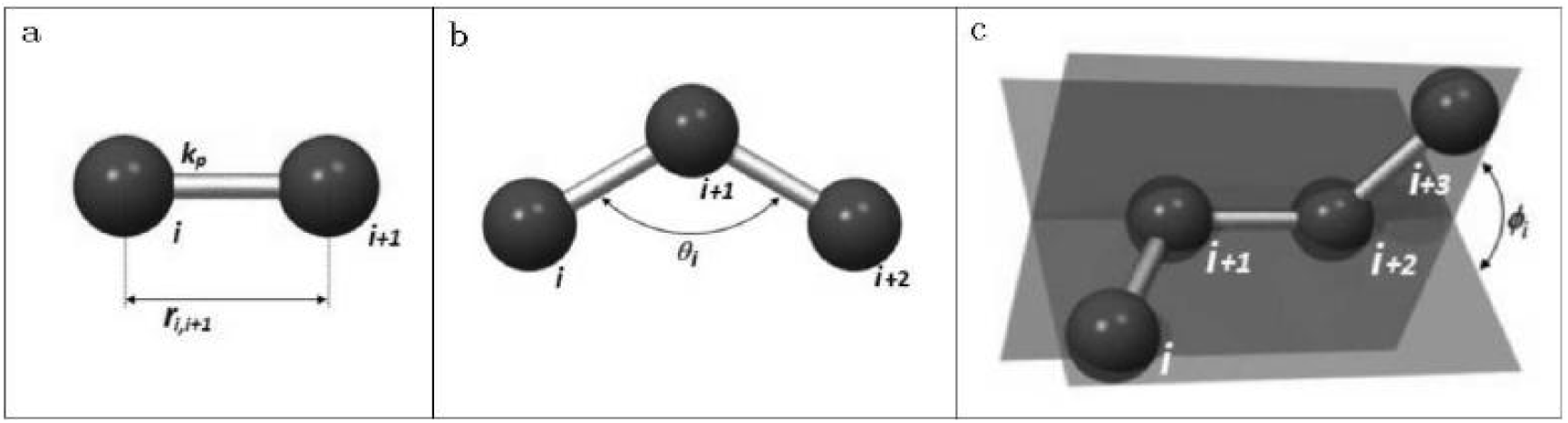
Panel a: Peptide bond schematic representation. Panel b: Sketch of the bending angle interaction. Panel c: Schematic view of the twist angle formed by two planes determined by four consecutive material points of the protein structure.

Let ***r***_*i*_(*i* = 1, …, *m*) be the position vector of the *m* monomer, and 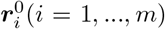 be the position vector of *m* residue in the current configuration. The peptide potential, responsible for the covalent bonds between the beads of the polymer chain, has the following expression:

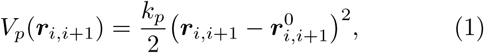

where 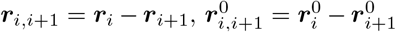 are the position vector between the bonded residues *i* and *i*+1 in the instantaneous and native configuration respectively. The norm of position vector is the bond length. The empirical constant 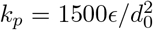 is in term of the equilibrium length parameter *d*_0_, and is in the unit of energy parameter *ϵ* In our simulation, *d*_0_ = 3:8 Å is the average distance between two monomers and *ϵ* sets the energy scale of the model. The bond potential is nothing more than a simple harmonic potential with spring constant *k_p_*.

Mathematically, the angular potential *V*_θ_ equivalent to peptide potential *V_p_* by replacing the relative displacement by angular difference, that is

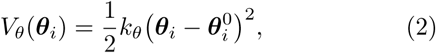

where *k_θ_* = 20**ϵ** rad^−2^ is the elastic constant expressed in term of the energy computational unit *ϵ*, and *θ_i_*, 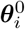are bond angles formed by three adjacent monomers in the simulated (time-dependent) and native conformation, respectively.

The dihedral potential (torsion) is 1-4 interaction, and are expressed as a function of the twisted angles *ϕ_i_* and 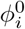, again refereed respectively to the actual and crystal configuration. The twisted angle is the angle formed between the two planes determined by four consecutive monomers along the chain. The definition of twisted angle potential *V_ϕ_* is

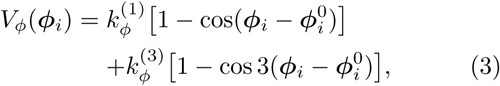

where 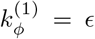 and 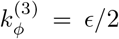 are dihedral constants expressed in term of energy unit *ϵ*.

Non-bonded (nb) interactions between nonconsecutive monomers are modeled with Lennard-John 12-10 potential. In Gō-like model a distinction is made among the pairs of residues that interact following a potential that has also an attractive part, in addition to a repulsive part. The criteria for this distinction is made on the basis of the native distance with respect to a parameter of the model, the so-called cut-off radius, *R_c_*. Native contact [78] are identified by the cut-off distance *R_c_* to be chosen such that two residues *i* and *j* form a native interaction if their distance *r_ij_* in the native state is less than cut-off distance *R_c_*. When two residues are not in native contact *r_ij_* > *R_c_*, they interact only through the Lennard-Jones repulsive tail (*σ/r_ij_*)^12^, where *σ* = 4.5 Å is a free length parameter correlated with the extension of the excluded volume (self-avoiding polymer chain). In other words, such residues in the polymer chain will interact attractively if brought to a distance greater than the native one and repulsive otherwise. We used *R_c_* = 3 Å in our simulation. The expression for Lennard-Jones potential is:

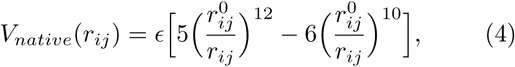

where all the symbols have been already defined. When 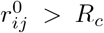, the purely repulsive contribution *V_nonnative_* is assigned to the pair of monomers considered. This term enhances the cooperativity in the folding process and takes the form of a Lennard-Jones barrier

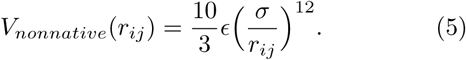

The non-bonded potential *V_nb_* summarized the possible long range interaction just described above and reads as

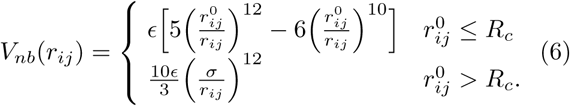

The total potential acting on all the residues of the polymer is then:

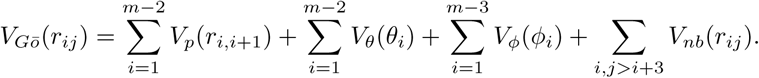

### B. Pore Model

The confinement effect on polymer dynamics can be represented by a step-like soft-core repulsive cylindrical potential. The cylinder axis of symmetry is set adjacent with the *x*-axis of the frame of reference used for polymer translocation simulation. The same *x*-axis direction is used to develop the mechanical pulling of the protein by using a constant force *F_x_* applied to the foremost beads inside the confinement. The constant force is used in analogy with the electrical potential in voltage driven translocation experiment. The schematic representation of the system, the potential and the pulling mechanism are shown in Fig. (2).

**FIG. 2.**
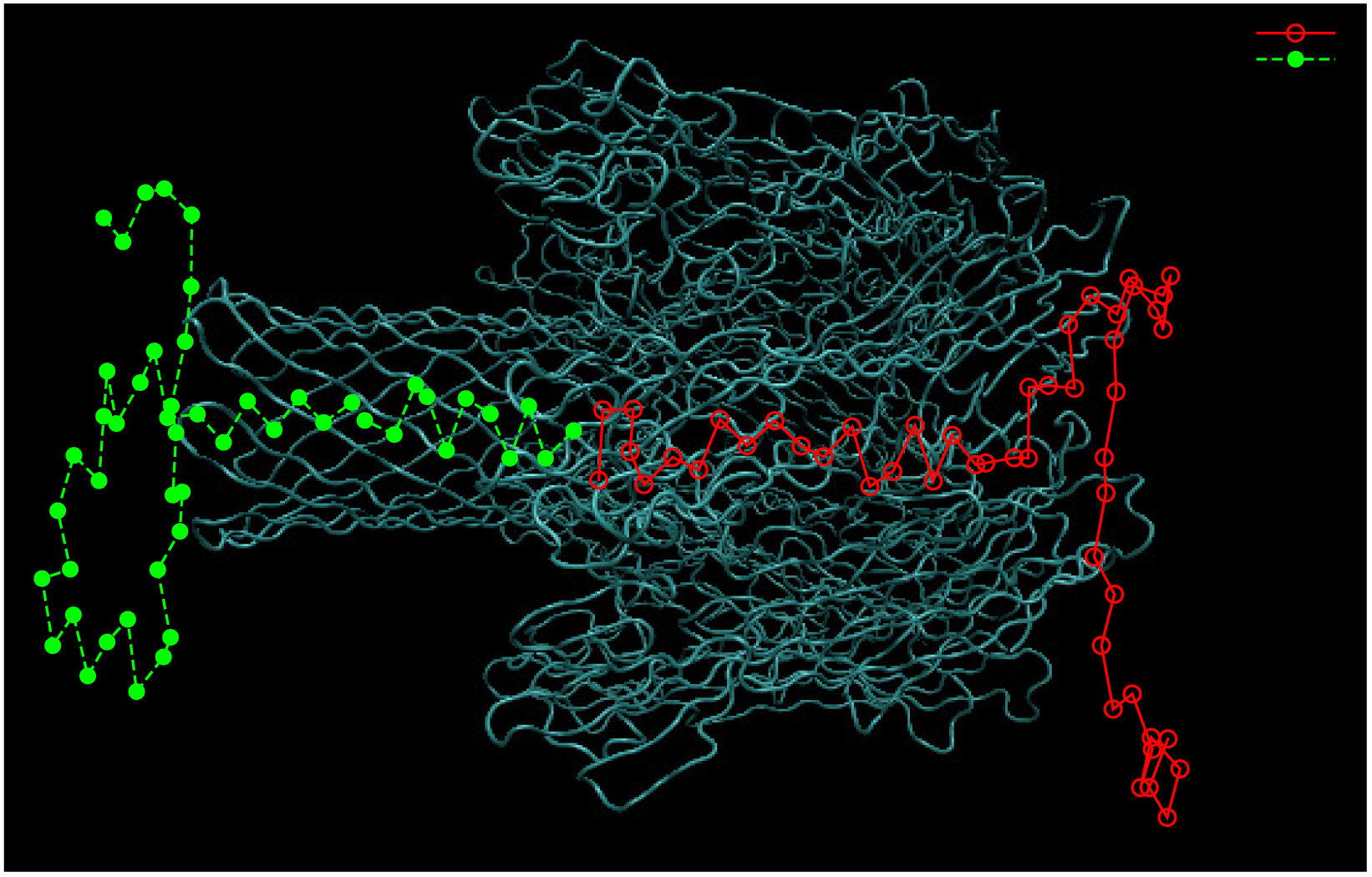
The confinement effect inside the pore is model by a step-like [0, *L*] cylindrical soft-core repulsive potential. A constant force *F_x_* in the direction of *x*-axis is used to drive the protein inside the channel. The force is always applied to the foremost residue inside the pore.

The expression of the pore potential is given by:

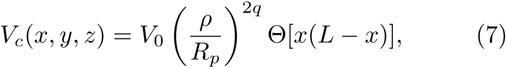

where *V*_0_ = 2*ϵ* and Θ(s) = [1 + tanh(αs)]/2 is a smooth step-like function limiting the action of the pore potential in the effective region [0;L]. *L* and *R*_p_ are pore length and radius respectively. Also, 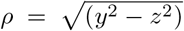 is the radial coordinate. The parameter *q* tunes the potential (soft-wall) stiffness, and α modulates the soft step-like profile in the *x*-direction; the larger the α, the steeper the step. In our simulation, we consider *q* = 1 and α = 2 Å^2^. The driving force *F_x_* acts only in the region in front of the pore mouth *x* ∈ [–2; 0], and inside the channel [0;L]. Pore length *L* = 100 Å and radius *R_p_* = 10 Å are taken from αHL structure data [79].

### C. Equation of Motion: Langevin dynamics

The equation of motion which governs the motion of material point is the well-know Langevin equation which is usually employed in coarse-grained molecular dynamics simulation. Consequently, numerical investigations are performed at constant temperature. The over-damped limit is actually implemented (*r¨* = 0) and a standard Verlet algorithm is used as numerical scheme for time integration [80]. The Langevin equation is given by

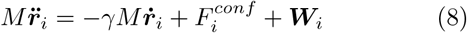

where 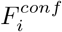 is the sum of all the internal and external forces acting on residue *i*. Here γ is the friction coefficient used to keep the temperature constant (also referred as Langevin thermostat). The random force ***W***_*i*_ accounts for thermal fluctuation, being a delta-correlated stationary and standard Gaussian process (white noise) with variance 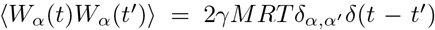. The random force satis_es the uctuation-dissipation theory; the mean-square of *W* is proportional to the corresponding friction coefficient γ.

## III. RESULTS

We used Langevin molecular dynamics simulation to transport polymer across a static *α*-hemolysin pore, as shown in Fig. (3). The total number of beads in the polymer is N = 50. We carry out Langevin molecular dynamics simulations using Verlet integration scheme [80]. The folding temperature occurred at *T ^∗^* = 0.77 in reduced temperature units *R*/*ϵ*, corresponding to the experimental denaturation temperature *T* = 338 K. This defined the energy scale to the value *ϵ* ≃0.88 kcal mol^*−*1^. Using the time scale *t_u_* = *σ*(*M*/120*ϵ*)^1*/*2^ we can obtained the physical time unit from the simulation time. With *ϵ* ≃ 0.88 kcal mol^*−*1^, *σ* = 4.5 Å, and assuming the average mass of monomer is *M* ~ 136 Da, we get *t_u_* ~ 0.25*ps*. In computer simulation the time step and friction coefficient used in the equation of motion (Langevin dynamics) are *h* = 0.001*t_u_* and 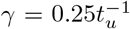, respectively. Further, the unit of force is defined as *f_u_* = *ϵ* Å^−1^ ~ 6 pN.

**FIG. 3.**
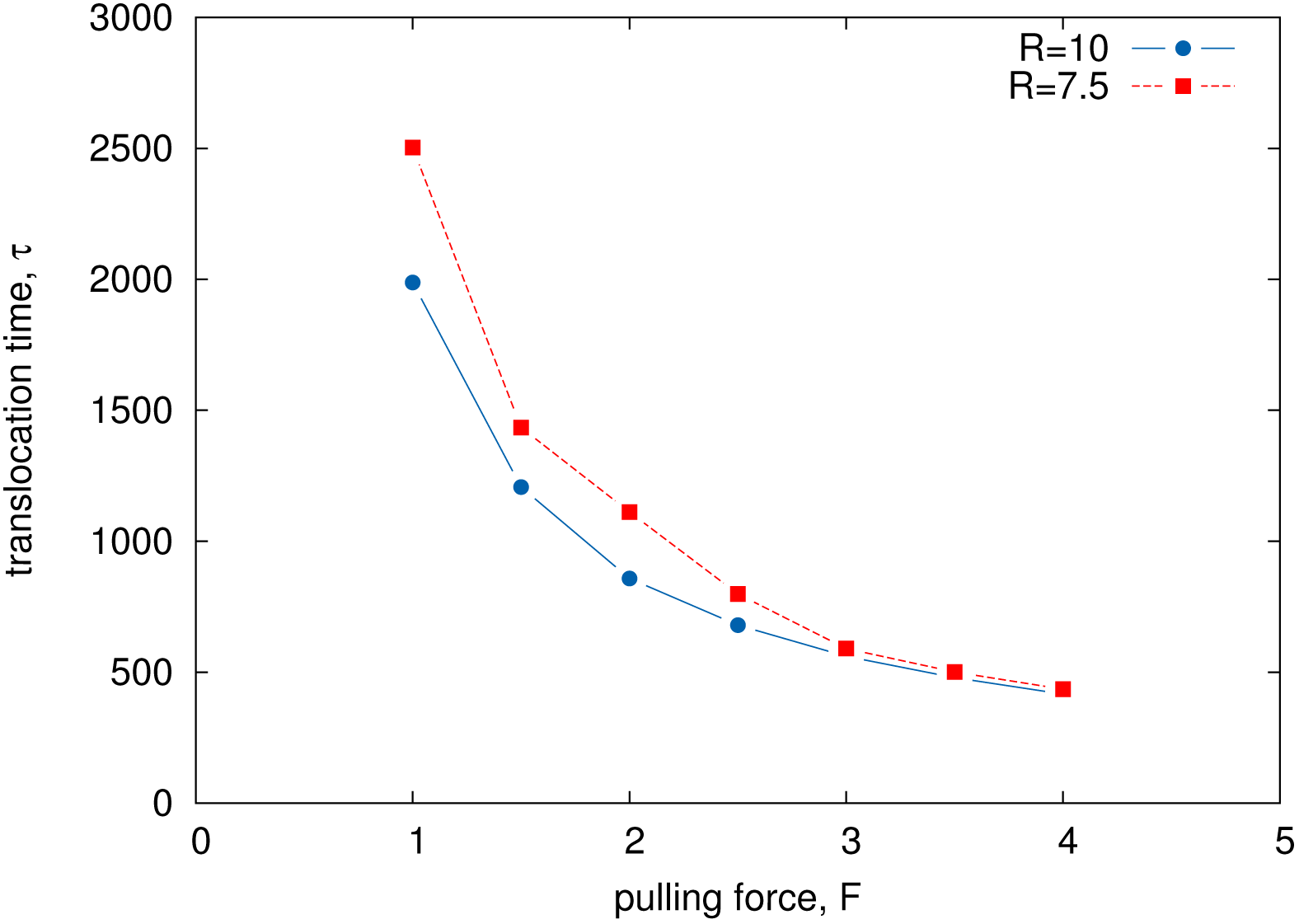
Conformation of polymer pulled across a *α*-hemolysin pore from CIS (left) to TRANS (right)-side, computed via a reduced-model simulation. The green solid circles represent the polymer on CIS-side moving towards the TRANS-side of the channel, while the red empty circle describe the polymer on the TRANS-side. The mechanical stretching reaction coordinate is the component of the end-to-end distance vector in the direction of the external force *F_x_* along the *x*-axis. The translocation coordinate is the displacement of the chain end along the axis of the cylindrical pore, relative to the pore entrance.

The simulation was run until the polymer was fully expelled out from the cis to tans-side. However, the translocation may fail at low external force *F* within a large assigned waiting time *t_w_*; in this case, the run stops, discards its statistics, and restarts a new trajectory. The probability of translocation *P_Tr_* can be defined as the number of translocation successes over the number of total runs, within the time *t_w_* ≃ 10^4^*t_u_*.

The effect of nanopore deformation play an important role on the translocation process. Recently, P. Fanzio, et al., [49] has shown that the DNA translocation process through an elastomeric nano-channel device can be altered by dynamically changing its cross section. In such experiment, the deformation in nano-channel is induced by a macroscopic mechanical compression of the polymer device. It has been demonstrated that it is possible to control and reversibly tune the direction of a nano-channel fabricated using elastomeric materials, so to fit the target molecule dimensions. The opportunity to dynamically control the nano-channel dimension open up new possibilities to understand the interactions between bio-molecules and nano-channels, such as the dependence of the translocation dynamics on the channel size, and the effects of moving walls [47]. Nano-pores embedded in thick synthetic membranes are also considered as barely deformable. Pore defromation properties may also play important role when tranlocating biological molecule (DNA, proteins) through graphene or other 2-dim crystals like MoS2.

Here we consider the situation where we change the radius of the pore from *R* = 10 to *R* = 7.5. We focus on the effect of the membrane deformability on the polymer translocation time *τ*. The translocation process can be characterized by studying the statistics of translocation times. The time statistics of translocation events is accessible to experiment [11] in which current loop drops signal the occupation of the channel by the passing molecule, for this reason, these times are also called the blockage times. In the simulation, the translocation time can be measured as first arrival time *t* at the channel end *x* = *L* of the protein center of mass. The average translocation time *τ* as a function of pulling force *F*, acting along the axis of the pore, are shown in figure (4). The blue line represent the average translocation time t when tranlocating polymer through α-hemolysin nanopore having radius *R* = 10 and length *L* = 100 compare with the result of small pore with radius *R* = 7:5, red dotted line in figure (4). At the high pulling force regimes, the average translocation time τ is not effected much. However, the translocation time increase at the low force regimes. As expected, stronger effects were observed for smaller pores when the pulling force is small. The average translocation time data points τ (*F*) as a function of pulling force F are consistent with an exponential behavior b (*F*)) exp(–*βF*). A decrease of the pore size implies the expected slowing-down of the average translocation time at fixed *F*. This effect, however, depends on the value of *F*, amounting to a modification of the overall shape of *τ* (*F*) mainly due to the presence of membrane-polymer interactions. From figure (4) it can be shown that only in the large force regime does it follow an Arrhenius-like law

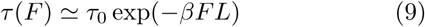

where *τ*_0_ being the Kramer’s time in the absence of the filed *F*, i.e *F* = 0, and *L* is the length of the pore.

**FIG. 4.**
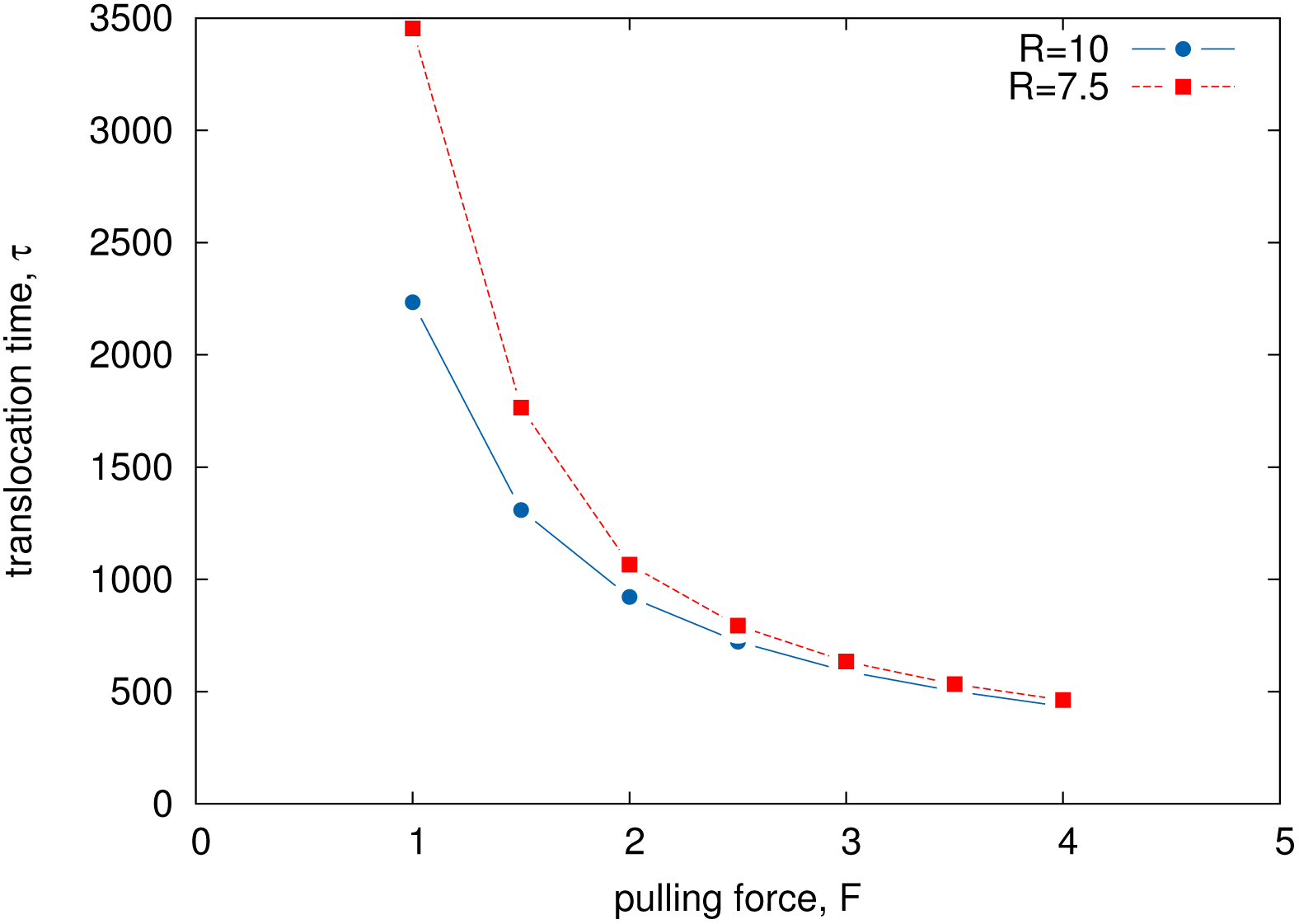
Average translocation times, *τ* with small (*R* = 7.5Å, red dashed line) and large (*R* = 10 Å, blue solid line) nanopores for the structured polymer of size N = 50, as a function of the pulling force *F*. Decreasing the size of the nano-pore slows down the translocation process at fixed *F*. At large *F* change are limited, whereas at low forces the size of the pore strongly influences the translocation. Here, we used 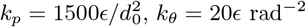, and dihedral constants 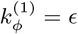 and 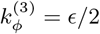.

To increase direct interaction between polymer and pore, we induce the deformation effect in polymer by changing the value of empirical constant *k_p_*, and dihedral constants 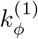 and 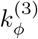, while keeping the elastic constant *k_θ_* unchanged, i.e. *k_θ_* = 20*ϵ* rad^−2^. Figure (5) illustrate the average translocation time as a function of pulling force for small pore of radius *R* = 7.5 and large pore of radius *R* = 10 with empirical constant *k_p_* = 3000*ϵ/d*^2^, dihedral constants 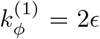 and 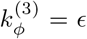. We observed significant increase in translocation time at the limit of low force for small pore. This clearly illustrate the strong interaction between the transport polymer and pore.

**FIG. 5.**
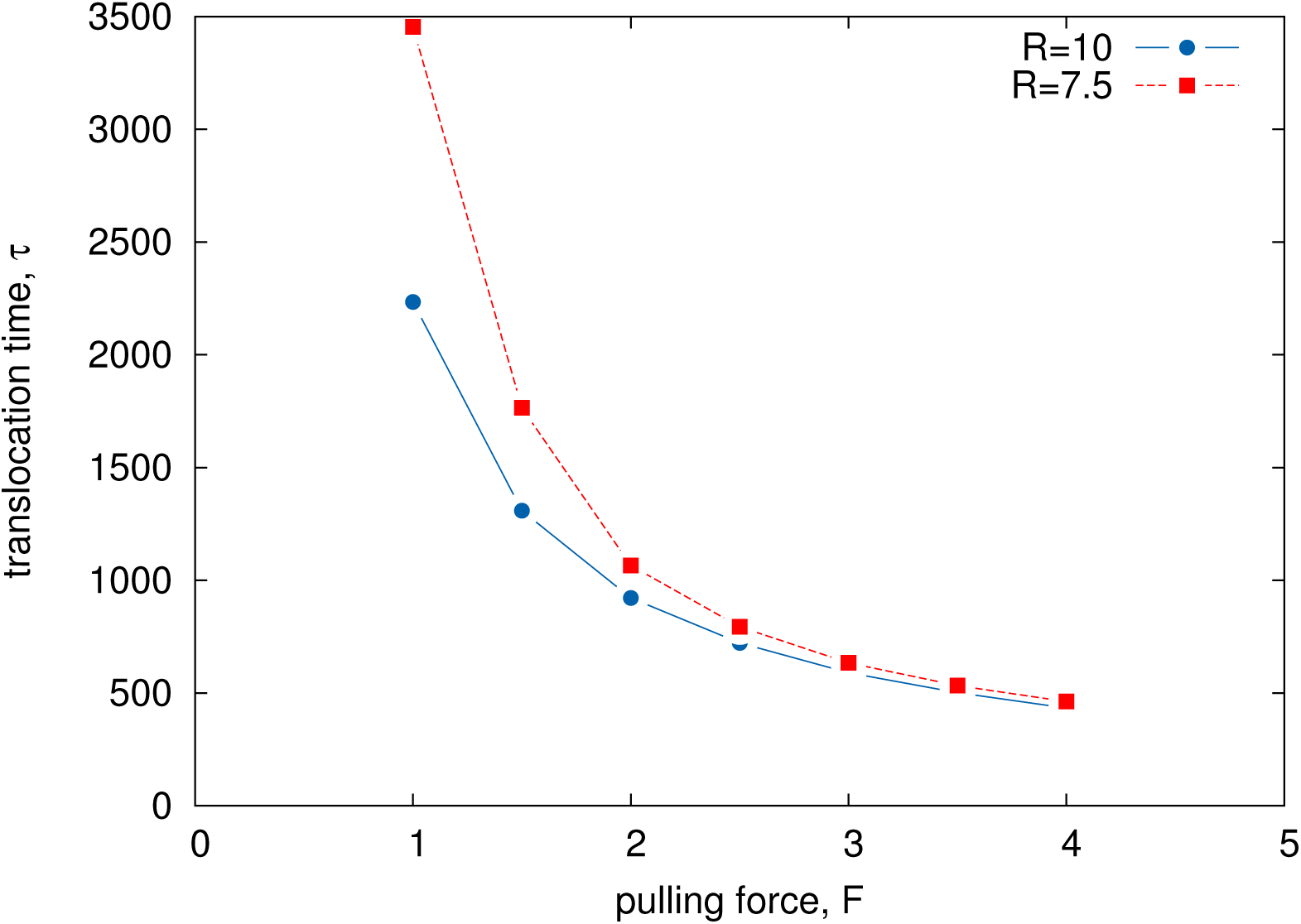
Average Translocation time *τ* as a function of force *F* for 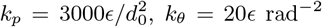, and dihedral constants 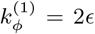 and 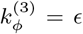. The red dashed line represent the time when polymer transport through a nanopore having radius *R* = 7.5 Å while the solid blue line describe the time with pore radius *R* = 10 Å.

**FIG. 6.**
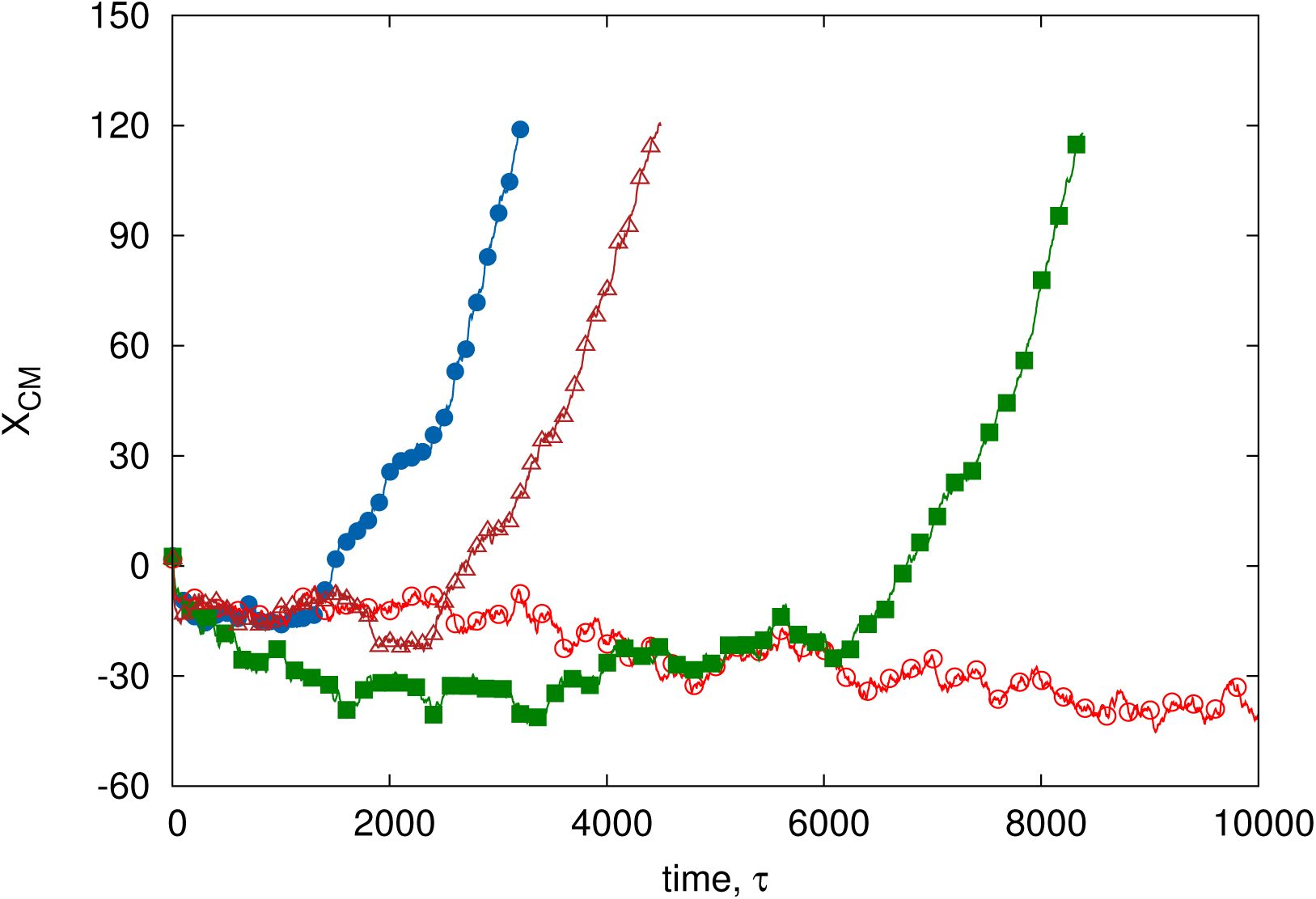
Trajectories of center of mass coordinate at the external force *F* = 1 and pore radius *R* = 10 for 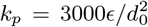 *k_θ_* = 20*ϵ* rad^−^2, and dihedral constants 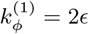 and 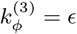. The red empty circle represent the fail translocation within the time window 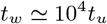.

Figure (**??**) describes the plot of typical unfolding trajectory as a function of time of GFP model when the force is applied to the C-terminal. Based on the topology map of GFP, an unfolding intermediate with the N-terminal residues still folded would require part of the beta-strand to be unstructured. As shown in Fig. (**??**), the folded intermediate stage enter as a loop inside the channel and translocates in double file conformation. Unfolding trajectory of the GFP molecule shows mainly one or two rips, each one followed by translocation of unfolded polypeptide chain [44].

Figure (7) shows the time distribution plot obtained by collecting the histogram from MD simulation in about 5000 numerical experiments of polymer translocation through a pore of radius *R* = 10 and length *L* = 1000 with pulling force *F* = 1 at temperature *T* = 198K. The distribution is not Gaussian as it presents a small degree of skewness. The green dotted line the fitting with Gaussian distribution. The analytical determination of the time distribution required the complete Smoluchowski problem to be solved (see appendix-A for more detail). Only approximately the data can be fitted via the function

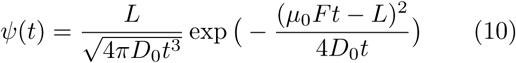

which is the distribution of first arrival time to the position *L* of biased random walkers starting from the origin in the assumptions of constant drift *v*_0_ = μ_0_*F*, semi-infinite channel [–∞*L*], and absorbing boundaries at *x* = –∞ and *L*.

**FIG. 7.**
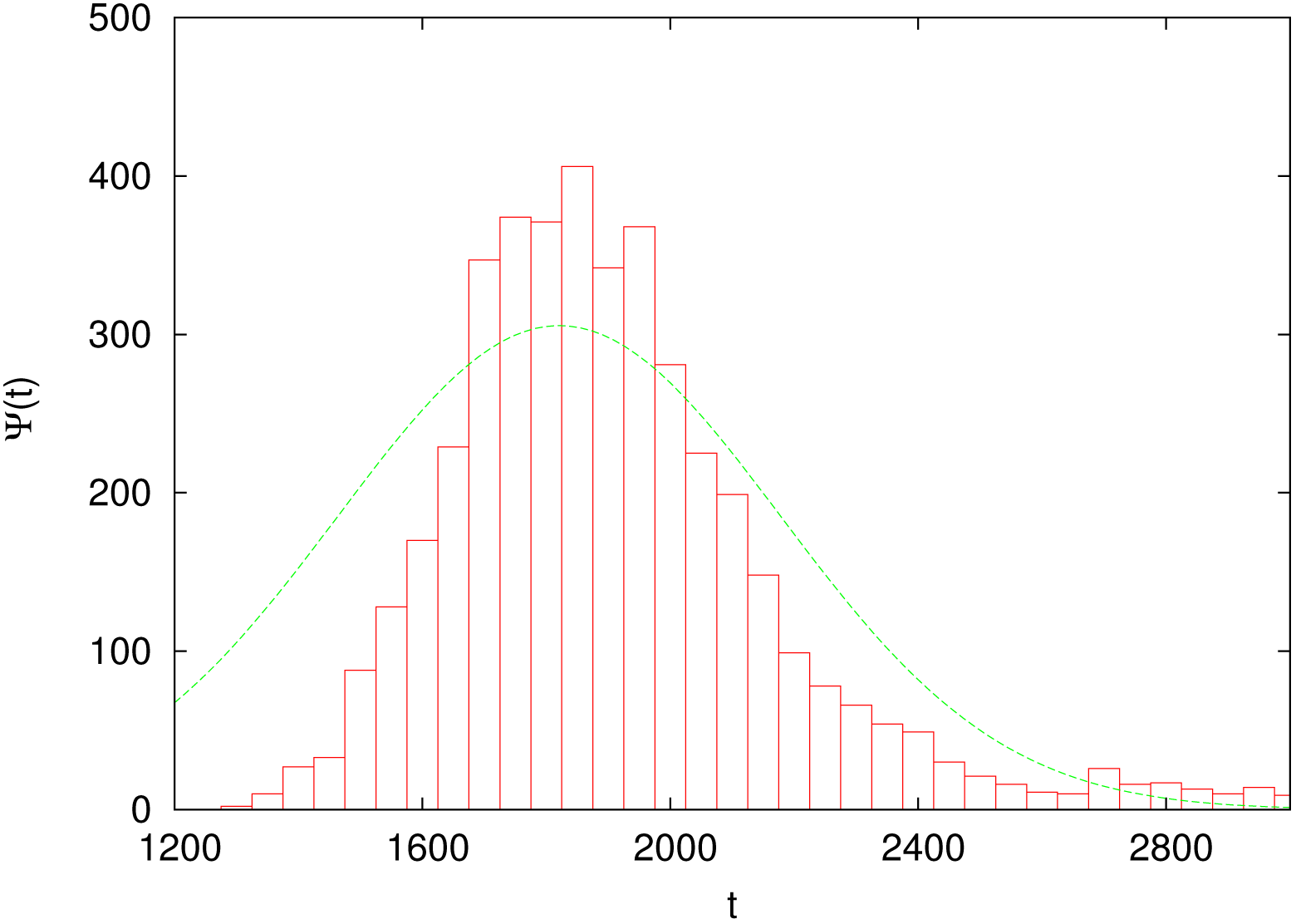
Distribution of translocation times across a static channel of radius *R_p_* = 10Å, and length *L* = 100 with 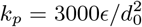, *k_θ_* = 20*ϵ* rad^−^2 and dihedral constants 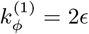 and 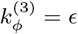 and force *F* = 1. The dashed line represents the fit via Gaussian

## IV. CONCLUSION

We have used coarse-grained computational model to simulate polymer translocation across *α*-hemolysin. In this simplified model translocation process we import a small polymer via a finite size channel by a uniform pulling force acting inside of the channel only. The molecular dynamics simulations at the coarse-grained level of protein translocation is compare to its theoretical interpretation by drift-diffusion Smoluchowski equation exploiting the first passage theory and radiation boundary condition. We examine the kinetic and thermodynamical characteristic of the process by studying the statistics of blockage times as a function of the pulling force *F* acting in the pore. As shown in the Fig. (4) and (5), the average translocation time translocation decrease when the importing force exceeds towards low limit. In contrast to large channel, the polymer takes longer time inside the channel and hence reveals strong interaction between polymer and pore. The results is further extended by varying the value of the bond and dihedral constant. Such deformation further increase the pore-polymer interaction and as a result average translocation time decrease at the limit of low pulling force. We also described the phenomenology of a coarse-grained computational model of polymer translocation process by the analytical formalism of the one-dimensional driven-diffusion Smoluchowski equation in the collective variable *X*. The theory reproduces the results of the average blockade times as a function of force *F*. Our theoretical approach contains almost all the information to describe the polymer translocation as a first-passage time problem of a driven diffusion stochastic process.

## ACKNOWLEDGMENTS

The author would like to thank U. M. B. Marconi, A. Vulpian and F. Cecconi for helpful discussion, and Institute of Complex Systems, National Research council of Rome for providing access to computational resources.

## Appendix A: Statistical Mechanics Description of Polymer Translocation

In order to interpret the numerical results it is appropriate to build suitable analytical models of the physical and chemical processes that occurred in real translocation and to define proper statistical observable quantities to grasp and resume the essence of the phenomenon. The translocation of a protein through a pore can be captured by a one-dimensional first-passage problem of a reaction coordinate *X*_1_, …., *X_k_* undergoing a driven-diffusion dynamics over a free-energy landscape. The dynamics of *X_j_* is stochastic in nature, so that we adopt a probabilistic description. We study the probability *P* (*X*_1_*, X_k_*; *t*) of visiting a state *X*_1_, …., *X_k_* at time, t, given an initial condition *P* (*X*_1_, ….*X_k_*; 0). We consider a long rod-like macromolecule of length *L* that translocates through a narrow channel in a thin membrane under the action of a constant force *F* [75, 77, 81–86, 88]. We are interested in a coarse-grained equation for the probability *P* (*x, t*) that a contour length *x* of the polymer’s chain has passed through the pore at time *t*. We assume that the polymer length *L* is much larger than the distance *d* between successive amino acids. For such a hydrodynamical description, we neglect the membrane thickness and describe the translocation process as diffusion of a point particle on an interval of length *L* in the presence of constant driving force *F*. Position of the particle on the interval at time *t* is equal to the length of that part of the macromolecule that has passed through the membrane by time *t*, as shown in Fig. (8). At time *t* = 0, the particle is injected onto the interval at *X* = 0. The equation of evolution for probability density *P* (*X*, *t*) of the reaction coordinate *X* can be described by effective Smoluchowski equation, defined as

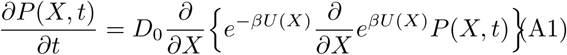

where, the potential *U*(*X*) = *G*(*X*) – *W*(*X*). Here, *G*(*X*) is the free-energy profile in the variable *X* (potential of mean force) of the unperturbed system and *W*(*X*) = *FX* is the work done on the system by the importing force *F*. Also in Eq. (A1), *D*_0_ is the effective diffusion related to effective mobility *μ*_0_ at equilibrium by the fluctuation dissipation relation: *μ*_0_ = *βD*_0_ and *β* = 1/(*k_B_T*). Through this way the phenomenology of translocation process can be conveniently recast as the motion of an effective particle of position *X* undergoing a driven diffusion in a potential *U*(*X*). Trans-location is considered completed when *X*(*t*) crosses a threshold value *X_th_*, corresponding to the protein outside the pore from the trans-side. This concept clearly defines a first-passage problem in a time window *T_w_* with exit time *t_out_* defined as *t_out_* = *min*{0 ≤*t* ≤*T_w_*|*X*(*t*) = *X_th_*}.

**FIG. 8.**
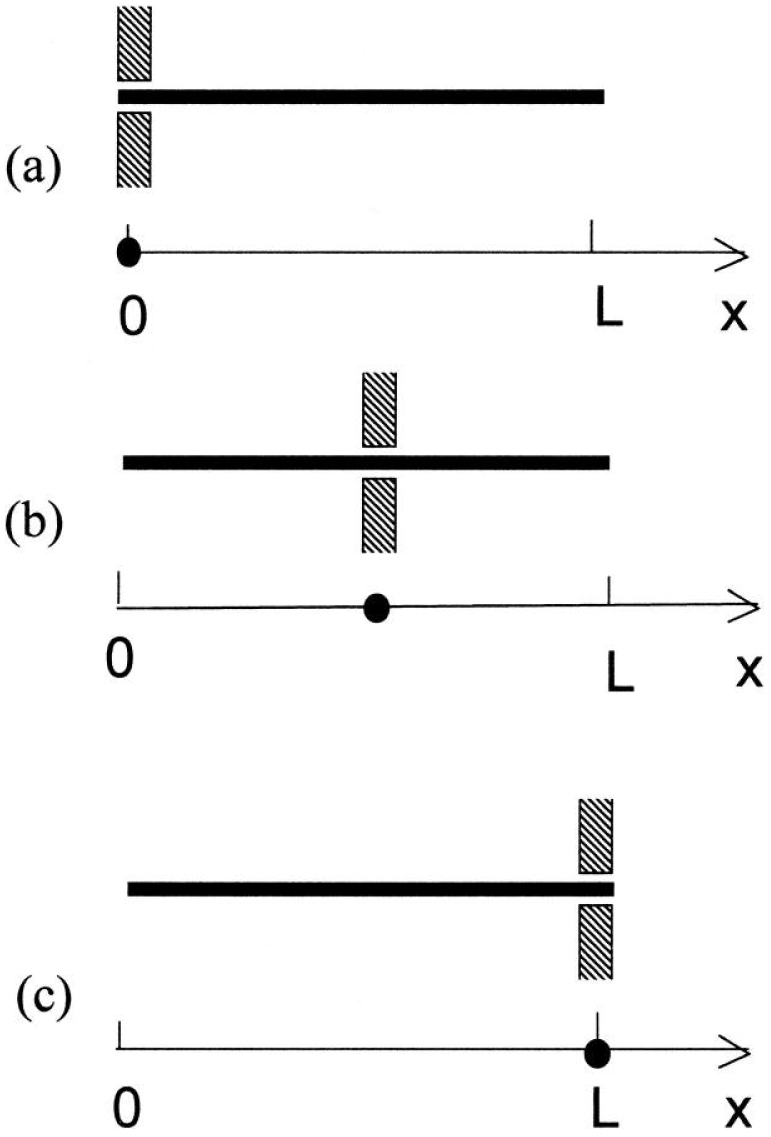
Schematic description of 1D diffusion model for translocation of a long rod-like macromolecule through a narrow channel in a thin membrane. Panel a: The entrance of the long rod-like macromolecule at *t* = 0. Panel b shows the molecule at an intermediate moment of time. Panel c shows the escape of molecule at *t* = 0.

Employing the radiative boundary condition for the current, we can write the expression for survival probability, that is, the probability that the molecule has not yet escaped the channel

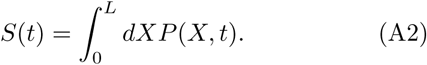

The survival probability *S*(*t*) estimates the number of systems whose *X*(*t*) is still in the interval [*X*_0_*, X_th_*], corresponding to the polymer still in channel, where *X*_0_ is the *X*-value at the pore entrance. The probability *S*(*t*) is related to the distribution *ψ*(*t*) of the exist time via the expression

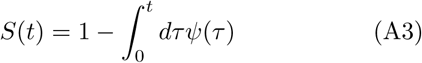

hence, by simple differentiation, the expression for translocation time distribution is

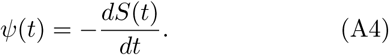

Using Eq. (A1) and (A2) and the boundary condition, the expression for *ψ*(*t*) becomes

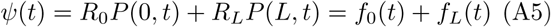

where *f*_0_(*t*) and *f_L_*(*t*) are the fluxes at the boundaries whose expression are obtained by solving the boundary value problem with the initial condition *P* (*X*, 0) = *δ*(*X* − *X*_0_). Where *R*_0_ and *R_L_* represents the rate at which the polymer expelled from the left and right side of channel, respectively.. The probability of translocation *P*_Tr_ can be defined by the integration of the flux *f_L_*(*t*) over all time, that is

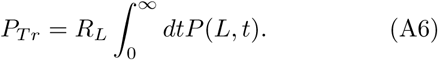

The average time spent in the channel by the molecule is given by

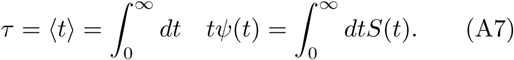

We can obtained approximate solution *P*(*Q*; *t*) of Eq. (A1) using Laplace transformation, define as

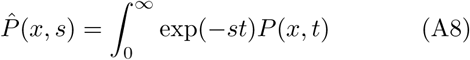

where *P*ˆ (*x*; *s*) is the Laplace transform of P(x; t). Using this, the partial differential Eq. (A1) becomes an ordinary differential equation

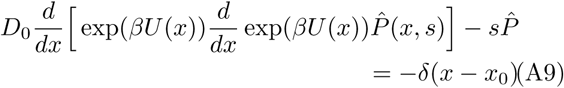

with the initial condition *P*(*x*; *t* = 0) = δ(*x* – *x*_0_).

The translocation probability and translocation time are obtained by the solution of Eq. (A9) for *s* = 0 and *Pˆ*(*x*; *s* = 0) (denoted by *P*ˆ (*x*) for the sake of simplicity).

Using Eq. (A6) and (A7), one can write

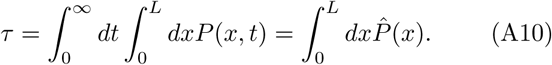

Introducing the function *Y* (*x*) = exp(*βU* (*x*))*P*ˆ (*x*), Eq. (A9) becomes

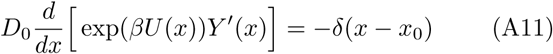

Using the theory of Green’s function, the solution of *Y* (*x*) of Eq. (A11) with boundary condition 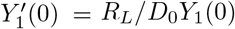 and 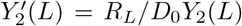 can be written as a superposition of two independent solution of *Y*_1_(*x*) and *Y*_2_(*x*) of the homogeneous equation. The two solution are

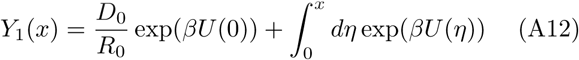

and

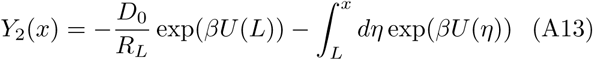

which provide the solution of Eq. (A9) for *s* = 0, in the form

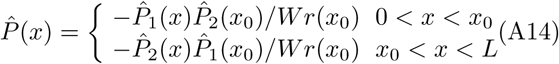

where the Wronskian 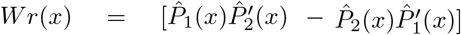 More detail on this method can be found in reference[77].

## References

[1] W. Wickner and R. Schekman. Protein translocation across biological membranes. Science, 310(5753), 1452:1456, 2005.

[2] M. T. Madigan, J. M. Matinko, and J. Parker. Biology of Micro-organisms. Prentice, Hall, Englewood Cliffs, NJ, 1997.

[3] B. Alberts, K. Robets, D. Bray, J. Lewis, M. Raff, and J. D. Watson. Molecular Biology of the Cell, Garland Publishing, New York and London, 1994.

[4] J.O. Bustamante, J. A. Hanover, and A. Liepins. The ion channel behavior of the nuclear pore complex. Journal of Membrane Biology, 146(3), 239:251, 1995.

[5] B. Hanss, E. Leal-Pinto, L. A. Bruggeman, T. D. Copeland, and P. E. Klotman. Identification and characterization of a cell membrane nucleic acid channel. Proceedings of the National Academy of Sciences, 95(4), 1921:1926, 1998.

[6] M.S. Sanford and Blobel G. A protein-conducting channel in the endoplasmic reticulum. Cell, 65(3), 371:380, 1991.

[7] A. Matouschek. Protein unfolding an important process in vivo? Opinion in Structural Biology, 13(1), 98:109, 2003.

[8] S. Prakash and A. Matouschek. Protein unfolding in the cell. Trends in Bio-chemical Sciences, 29(11), 593:600, 2004.

[9] M.S. Sanford and Blobel G. A protein-conducting channel in the endoplasmic reticulum. Cell, 65(3), 371:380, 1991.

[10] John J. Kasianowicz, Eric Brandin, Daniel Branton, and David W. Deamer. Characterization of individual polynucleotide molecules using a membrane channel. Proc. Natl. Acad. Sci. USA, 93, 13770:13773, 1996.

[11] Amit Meller, Lucas Nivon, and Daniel Branton. Voltage-Driven DNA Translocations through a Nanopore. Phys. Rev. Lett., 86, 3435, 2001.

[12] John J. Kasianowicz, Eric Brandin, Daniel Branton, and David W. Deamer. Characterization of individual polynucleotide molecules using a membrane channel. Proc. Natl. Acad. Sci. USA, 93, 13770:13773, 1996.

[13] Amit Meller, Lucas Nivon, Eric Brandin, Jene Golovchenko, and Daniel Branton. Rapid nanopore discrimination between single polynucleotide molecules. Biochemistry, 97, 1079:1084, 1999.

[14] Jerome Mathe, Aleksei Aksimentiev, David R. Nelson, Klaus Schulten, and Amit Meller. Orientation discrimination of single-stranded DNA inside the *α*-hemolysin membrane channel. Proc. Natl. Acad. Sci. USA, 102, 12377:12382, 2005.

[15] Ryuji Kawano, Anna E. P. Schibel, Christopher Cauley and Henry S. White. Controlling the Translocation of Single-Stranded DNA through-Hemolysin Ion Channels Using Viscosity. Langmuir, 25, 1233:1237, 2009

[16] Alexis F. Sauer-Budge, Jacqueline A. Nyamwanda, David K. Lubensky, and Daniel Branton. Unzipping Kinetics of Double-Stranded DNA in a Nanopore. Phys. Rev. Lett., 90, 238101, 2003.

[17] M. Bates, M. Burns, A. Meller. Dynamics of DNA molecules in a membrane channel probed by active control techniques. Biophys J., 84, 2366:72, 2003.

[18] Jeff Nivala, Douglas B. Marks, and Mark Akeson. Unfoldase-mediated protein translocation through an *α*- hemolysin nanopore. Nature Biotechnology, 31, 247:250, 1013

[19] M. M. Mohammad, S. Prakash, A. Matouschek. and L. Movileanu. Controlling a single protein in a nanopore through electrostatic traps. J. Am. Chem. Soc., 130, 4081:4088, 2008.

[20] Liu, J. et al. Polarization-induced local pore-wall functionalization for biosensing: From micropore to nanopore. Analytical Chemistry 84, 32543261, 2012.

[21] R. S. Wei, V. Gatterdam, R. Wieneke, R. Tampe, U. Rant. Stochastic sensing of proteins with receptor-modified solid-state nanopores. Nature Nanotechnology, 7, 257:263, 2012.

[22] D. S. Talaga and J. L. Li . Single-Molecule Protein Unfolding in Solid State Nanopores. Journal of the American Chemical Society, 131, 9287:9297, 2009.

[23] S. W. Kowalczyk, A. R. Hall and C. Dekker. Detection of Local Protein Structures along DNA Using Solid-State Nanopores. Nano Letters, 10, 324:328, 2010.

[24] W. J. Lan, D. A. Holden, B. Zhang, H. S. White. Nanoparticle Transport in Conical-Shaped Nanopores. Analytical Chemistry, 83, 3840:3847, 2011.

[25] A. R. Hall, J. M. Keegstra, M. C. Duch, M. C. Hersam, C. Dekker. Translocation of Single-Wall Carbon Nanotubes Through Solid-State Nanopores. Nano Letters, 11, 2446:2450, 2011.

[26] Q. Liu, Wu H. Wu L, Xie X, Kong J, et al. Voltage-Driven Translocation of DNA through a High Throughput Conical Solid-State Nanopore. PLoS ONE, 7(9), e46014, 2012.

[27] M. Wanunu, Sutin J, McNally B, Chow A, Meller A. DNA Translocation Governed by Interactions with Solid-State Nanopores. Biophysical Journal, 95, 4716:4725, 2008.

[28] Arnold J. Storm, Cornelis Storm, Jianghua Chen, Henny Zandbergen, Jean-Franois Joanny, and Cees Dekker. Fast DNA Translocation through a Solid-State Nanopore. Nano Lett., 5, 1193:1197, 2005.

[29] Breton Hornblower, Amy Coombs, Richard D Whitaker, Anatoly Kolomeisky, Stephen J Picone, Amit Meller and Mark Akeson. Single-molecule analysis of DNA-protein complexes using nanopores. Nature Methods, 4, 315:317, 2007.

[30] Piere Rodriguez-Aliaga, Luis Ramirez, Frank Kim, Carlos Bustamante and Andreas Martin. Substrate-translocating loops regulate mechanochemical coupling and power production in AAA+ protease ClpXP. Nature Structural and Molecular Biology, 23, 974–981, 2016.

[31] E. S. Johnson, I. Schwienhorst, R. J. Dohmen, and G. Blobel. The ubiquitin-like protein Smt3p is activated for conjugation to other proteins by an Aos1p/Uba2p heterodimer. EMBO J., 16, 5509:5519, 1997.

[32] T. A. Baker, and Sauer, R.T. ClpXP, an ATP-powered unfolding and protein-degradation machine. Biochim. Biophys. Acta, 1823, 15:28, 2012.

[33] Aubin-Tam, M.E. et al. Single-molecule protein unfolding and translocation by an ATP-fueled proteolytic machine. Cell, 145, 257:267, 2011.

[34] S. Gottesman, Roche, E., Zhou, Y. and Sauer, R.T. The ClpXP and ClpAP proteases degrade proteins with carboxy-terminal peptide tails added by the SsrA-tagging system. Genes Dev., 12, 1338:1347, 1998.

[35] Y. Kim, et al. Dynamics of substrate denaturation and translocation by the ClpXP degradation machine. Mol. Cell, 5, 639:648, 2000.

[36] A. Martin, Baker, T.A. and Sauer, R.T. Rebuilt AAA+ motors reveal operating principles for ATP-fuelled machines. Nature, 437, 1115:1120, 2005.

[37] Maya Sen, Rodrigo A. Maillard, Kristofor Nyquist, Piere Rodriguez-Aliaga, Steve Press, Andreas Martin, Carlos Bustamante. The ClpXP Protease Unfolds Substrates Using a Constant Rate of Pulling but Different Gears. Cell, 155, 636:646, 2013.

[38] A. Kravats, Jayasinghe M, Stan G. Unfolding and translocation pathway of substrate protein controlled by structure in repetitive allosteric cycles of the ClpY ATPase. Proc Natl Acad Sci U S A., 108, 2234:9, 2011.

[39] Nobuyasu Koga, Tomoshi Kamedac, Kei-ichi Okazakib and Shoji Takada. Paddling mechanism for the substrate translocation by AAA+ motor revealed by multiscale molecular simulations. Proc Natl Acad Sci U S A., 106, 18237:18242, 2009.

[40] Y. Shin, Davis J. H, Brau R. R, Martin A, Kenniston J.A, Baker T. A, Sauer R.T, Lang M. J. Single-molecule denaturation and degradation of proteins by the AAA+ ClpXP protease. Proc Natl Acad Sci U S A., 106, 19340:5. 2009.

[41] R. A. Maillard, et al. ClpX(P) generates mechanical force to unfold and translocate its protein substrates. Cell, 145, 459:469, 2011.

[42] Hendrik Dietz and Matthias Rief. Exploring the energy landscape of GFP by single-molecule mechanical experiments. Proc Natl Acad Sci U S A., 101, 16192:16197, 2004.

[43] Moritz Mickler, Ruxandra I. Dima, Hendrik Dietz, Changbong Hyeon, D. Thirumalai, and Matthias Rief. Revealing the bifurcation in the unfolding pathways of GFP by using single-molecule experiments and simulations. Proc Natl Acad Sci U S A., 104, 20268:20273, 2007.

[44] R. A. Maillard, et al. ClpX(P) generates mechanical force to unfold and translocate its protein substrates. Cell, 145, 459:469, 2011.

[45] Hendrik Dietz and Matthias Rief. Exploring the energy landscape of GFP by single-molecule mechanical experiments. Proc Natl Acad Sci U S A., 101, 16192:16197, 2004.

[46] Moritz Mickler, Ruxandra I. Dima, Hendrik Dietz, Changbong Hyeon, D. Thirumalai, and Matthias Rief. Revealing the bifurcation in the unfolding pathways of GFP by using single-molecule experiments and simulations. Proc Natl Acad Sci U S A., 104, 20268:20273, 2007.

[47] J. A. Cohen, Chaudhuri, A. and Golestanian R. Active Polymer Translocation through Flickering Pores. Phys. Rev. Lett., 107, 238102, 2011.

[48] H. Zhang, Zhao Q, Tang Z, Liu S, Li Q, Fan Z, Yang F, You L, Li X, Zhang J, Yu D. Slowing down DNA translocation through solid-state nanopores by pressure. Small., 9, 4112:7, 2013.

[49] Paola Fanzio, Chiara Manneschi, Elena Angeli, Valentina Mussi, Giuseppe Firpo, Luca Ceseracciu, Luca Repetto and Ugo Valbusa. Modulating DNA Translocation by a Controlled Deformation of a PDMS Nano-channel Device. Scientific Reports, 2, 791, 2012.

[50] D. Huh, Mills KL, Zhu X, Burns MA, Thouless MD, Takayama S. Tuneable elastomeric nano-channels for nanofluidic manipulation. Nat Mater., 6, 424:8, 2007.

[51] E. Angeli, Manneschi C., Repetto L., Firpo G. and Valbusa U. DNA Manipulation with Elastomeric Nanostructures Fabricated by Soft-Moulding of a FIB-Patterned Stamp. Lab Chip, 11, 2625, 2011.

[52] Nicholas A. W. Bell, Murugappan Muthukumar, Ulrich F. Keyser Translocation frequency of double-stranded DNA through a solid-state nanopore. arXiv:1508.04396 [physics.bio-ph], 2015.

[53] T. Ikonen, J. Shin, W. Sung, T. Ala-Nissila. Polymer translocation under time-dependent driving forces: Resonant activation induced by attractive polymer-pore interactions. The Journal of chemical physics, 136, 205104, 2012.

[54] A. Fiasconaro, J. J. Mazo, and F. Falo. Active polymer translocation in the three-dimensional domain. Phys. Rev. E, 91, 022113, 2015.

[55] Stefureac R. I, Kachayev A, Lee J. S. Modulation of the translocation of peptides through nanopores by the application of an AC electric field. Chem. Commun., 48, 1928:30, 2012.

[56] M. Bates, Burns M, Meller A. Dynamics of DNA molecules in a membrane channel probed by active control techniques. Biophys J., 84, 2366:72, 2003.

[57] Jalal Sarabadani, Timo Ikonen, Tapio Ala-Nissila Theory of polymer translocation through a flickering nanopore under an alternating driving force. arXiv:1505.04057 [cond-mat-statmech]., 2015.

[58] Piotr Szymczak. Periodic forces trigger knot untying during translocation of knotted proteins. Scientific Reports, 6, 21702, 2016.

[59] J. Yamada, J. L. Phillips, S. Patel et al., A bimodal distribution of two distinct categories of intrinsically disordered structures with separate functions in FG nucleoporins. Mol Cell Proteomics., 9, 2205:24, 2010.

[60] P. Rehling, Brandner K, Pfanner N. Mitochondrial import and the twin-pore translocase. Nat. Rev. Mol. Cell Biol., 5, 519:30, 2004.

[61] G. Sigalov, Jeffrey Comer, Gregory Timp, and Aleksei Aksimentiev. Detection of DNA sequences using an alternating electric field in a nanopore capacitor. Nano Lett., 8, 56:63, 2008.

[62] Gregory F Schneider and Cees Dekker. DNA sequencing with nanopores. Nature Biotechnology, 30, 326:328, 2012.

[63] Meni Wanunu. Nanopores: A journey towards DNA sequencing. Phys Life Rev. 9, 125:158, 2012.

[64] Timothée Menais, Stefano Mossa and Arnaud Buhot. Polymer translocation through nano-pores in vibrating thin membranes. Scientific Reports, 6, 38558, 2016.

[65] Fabio Cecconi, Muhammad Adnan Shahzad, Umberto Marini Bettolo Marconi and Angelo Vulpiani. Frequency-control of protein translocation across an oscillating nanopore. Phys. Chem. Chem. Phys., 19, 11260–11272, 2017.

[66] N. Gō and H. A. Scheraga. On the use of classical statistical mechanics in the treatment of polymer chain conformations. Macromolecules, 9, 535:542, 1976.

[67] C. Clementi, H. Nymeyer, and J. N. Onuchic. Topological and energetic factors: what determines the structural details of the transition state ensemble and on- route intermediates for protein folding? An investigation for small globular proteins. J. Mol. Biol., 298, 937:953, 2000.

[68] N. Gō. Theoretical studies of protein folding. Ann. Rev. of Biophys. and Bioeng., 12, 183, 1983.

[69] E. Shakhnovich. Theoretical studies of protein-folding thermodynamics and kinetics. Current Opinion in Structural Biology, 7, 29:40, 1997.

[70] D. Baker. A surprising simplicity to protein folding. Nature, 405, 39:42, 2000.

[71] J.K. Karanicolas and C. L. Brooks. Improved gō-like models demonstrate the robustness of protein folding mechanisms towards non-native interactions. J. Mol. Bio., 334, 309:325, 2003.

[72] J.N. Onuchic and P. G. Wolynes. Theory of protein unfolding. Curr. Opin. Struct. Bio., 14, 70:75, 2004.

[73] R.D. Jr. Hills and C. L. Brooks. Insights from coarse-grained gō-models for protein folding and dynamics. Int. J. of Mol. Sci., 10, 889:905, 2009.

[74] C. Guardiani, M. Cencini, F. Cecconi. Coarse-grained modeling of protein unspecifically bound to DNA. Phys. Bio., 11, 026003 2014.

[75] F. Cecconi, M. Bacci, M. Chinappi. Protein transport across nanopores: a statistical mechanical perspective from coarse-grained modeling and approaches. Prot. Pept. Lett., 21, 227:234 2014.

[76] M. Chinappi, F. Cecconi, C. M. Casciola. Computational analysis of Maltose Binding Protein translocation. Philos. Mag., 91, 2034:2048 2011.

[77] A. Ammenti, F. Cecconi, U. Marini-Bettolo-Marconi and A. Vulpiani. A statistical model for translocation of structured polypeptide chains through nanopores. J. Phys. Chem. B, 113, 10348 2009.

[78] S. Miyazawa, Jernigan R. L. Residue-residue potentials with a favorable contact pair term and an unfavorable high packing density term, for simulation and threading. J Mol Biol., 256, 623:44, 1996.

[79] M. Chinappi, F. Cecconi, and C.M. Casciola. Computational analysis of maltose binding protein translocation. Philos. Mag., 91, 2034:2048, 2011.

[80] L. Verlet. Computer experiments on classical fluids. i. thermodynamical properties of Lennard-Jones molecules. Phys. Rev., 159, 98:103, 1967.

[81] A. M. Berezhkovskii, M. A. Pustovoit, and S. M. Bezrukov. Channel-facilitated membrane transport: Transit probability and interaction with the channel. Journal of Chemical Physics, 116, 9952:9956, 2002.

[82] A. M. Berezhkovskii and I. V. Gopich. Translocation of Rod-like Polymers through Membrane Channels. Biophys J., 84, 787:793, 2003.

[83] D. K. Lubensky, and D. R. Nelson. Driven polymer translocation through a narrow pore. Biophys. J., 77, 1824:1838, 1999.

[84] N. Lee and and S. Obukhov. Diffusion of a polymer chain through a thin membrane. J. Phys. II., 6, 195:204, 1996.

[85] Muthukumar, M. Polymer translocation through a hole. J. Chem. Phys., 111, 10371:10374, 1999.

[86] Muthukumar, M. Translocation of a confined polymer through a hole. Phys. Rev. Lett., 86, 3188:3191, 2001

[87] P. P. Valko and S. Vajda. Inversion of noise-free Laplace transforms: towards a standardized set of test problems. Inverse Problems in Engineering., 10, 467:483, 2002.

[88] P. Ansalone, M. Chinappi, L. Rondoni, F. Cecconi. Driven diffusion against electrostatic or effective energy barrier across-Hemolysin. J. Chem. Phys., 143, 154109 2015.

